# Accuracies of genomic predictions for disease resistance of striped catfish to *Edwardsiella ictaluri* using artificial intelligence algorithms

**DOI:** 10.1101/2021.05.10.443499

**Authors:** Nguyen Thanh Vu, Tran Huu Phuc, Kim Thi Phuong Oanh, Nguyen Van Sang, Trinh Thi Trang, Nguyen Hong Nguyen

**Author notes:** Correspondence to N.T. Vu or and N.H. Nguyen.

## Abstract

Assessments of genomic prediction accuracies using artificial intelligence (AI) algorithms (i.e., machine and deep learning methods) are currently not available or very limited in aquaculture species. The principal aim of this study was to examine the predictive performance of these new methods for disease resistance to *Edwardsiella ictaluri* in a population of striped catfish *Pangasianodon hypophthalmus* and to make comparisons with four common methods, i.e., pedigree-based best linear unbiased prediction (PBLUP), genomic-based best linear unbiased prediction (GBLUP), single-step GBLUP (ssGBLUP) and a non-linear Bayesian approach (notably BayesR). Our analyses using machine learning (i.e., ML-KAML) and deep learning (i.e., DL-MLP and DL-CNN) together with the four common methods (PBLUP, GBLUP, ssGBLUP and BayesR) were conducted for two main disease resistance traits (i.e., survival status coded as 0 and 1 and survival time, i.e., days that the animals were still alive after the challenge test) in a pedigree consisting of 560 individual animals (490 offspring and 70 parents) genotyped for 14,154 Single Nucleotide Polymorphism (SNPs). The results using 6470 SNPs after quality control showed that AI methods outperformed PBLUP, GBLUP and ssGBLUP, with the increases in the prediction accuracies for both traits by 9.1 – 15.4%. However, the prediction accuracies obtained from AI methods were comparable to those estimated using BayesR. Imputation of missing genotypes using AlphaFamImpute increased the prediction accuracies by 5.3 – 19.2% in all the methods and data used. On the other hand, there were insignificant decreases (0.3 – 5.6%) in the prediction accuracies for both survival status and survival time when multivariate models were used in comparison to univariate analyses. Interestingly, the genomic prediction accuracies based on only highly significant SNPs (P < 0.00001, 318 - 400 SNPs for survival status and 1362 – 1589 SNPs for survival time) were somewhat lower (0.3 to 15.6%) than those obtained from the whole set of 6,470 SNPs. In most of our analyses, the accuracies of genomic prediction were somewhat higher for survival time than survival status (0/1 data). It is concluded that there are prospects for the application of genomic selection to increase disease resistance to *Edwardsiella ictaluri* in striped catfish breeding programs.

## 1. Introduction

Genomic selection has been increasingly practiced in genetic improvement programs for farmed animals and plants, using a range of different statistical methods from genomic-based best linear unbiased prediction (GBLUP) and its extension known as single-step GBLUP (ssGBLUP) to non-linear Bayesian approaches (BayesA, BayesB, BayesC, BayesC-*π* and notably BayesR) (Lourenco et al. 2020; Moser et al. 2015; VanRaden 2008; Daetwyler et al. 2013). Recently, there has been a growing interest in using Artificial Intelligent (AI) algorithms (i.e., Machine Learning or Deep Learning) to choose optimal genome-wide models without prior consumptions to determine genomic prediction accuracies for quantitative complex traits, especially for disease resistance (tolerance or resilience). Examples using simulated and real animal and plant data showed that the prediction accuracies (*r*) using machine learning were 10% greater than linear (i.e. GBLUP) and 1.3% greater than non-linear methods (i.e. BayesR), for instance, *r* = 0.732 – 0.758 for binary and continuous traits using GBLUP vs 0.801 – 0.832 (Yin et al. 2020) or the increase of 20 – 70% in accuracy when using machine learning (i.e., linear bagging) for disease resistance to photobacterium in gilthead sea bream (Bargelloni et al. 2021). To date, however, no (or very limited) studies have used simultaneously AI methods to estimate genomic breeding values for aquaculture species.

Almost all studies in aquatic animal species have employed GBLUP, ssGBLUP or Bayesian methods to examine genomic prediction accuracies for two main groups of traits, i.e., body weight and disease resistance. A synthesis of the published information shows that the genomic prediction accuracy ranged from 0.38 to 0.89 for growth-related traits (Houston et al. 2020). On the other hand, the predictions of genomic breeding values for disease resistance were varied with studies, pathogens, populations and methods used, with the estimates ranging from 0 (Silva et al. 2019) to 0.8 or greater (Barría et al. 2018). For both growth and disease traits, the majority of the studies demonstrated that either linear (GBLUP, ssGBLUP) or non-linear Bayesian methods increased the prediction accuracies of animal breeding values relative to Pedigreed-based BLUP (PBLUP) by 22 - 24% (Houston et al. 2020). In addition to growth and disease traits, three recent studies performed genomic predictions for meat quality in banana shrimp (Nguyen et al. 2020) or Portuguese Oyster (*Crassostrea angulata*) (Vu et al. 2021) as well as behavior traits (i.e., cannibalism) in Asian seabass (Nguyen and Khang 2021).

Despite the economic importance of the striped catfish in aquaculture, e.g., valuing 2.36 billion USD that accounts for 1% GDP of Vietnam, there has been a paucity of knowledge in using genomic information in breeding programs for high growth (Vu et al. 2019b) or increased disease resistance, especially Bacillary Necrosis of Pangasius (Vu et al. 2019a). The Bacillary Necrosis of Pangasius (BNP) due to *Edwardsiella ictaluri* has been the main contributor to 56 – 92% mortality during larval and fingerling rearing in this species (Vũ et al. 2017). Our earlier studies showed that there is heritable genetic component for *E. ictaluri* resistance, but the heritability for this trait was low, around 0.10 across statistical methods used (Vu et al. 2019a; Pham et al. 2021; Dinh Pham et al. 2021). While these results suggests there are prospects for genetic improvement of resistance to *E. ictaluri* using conventional selective breeding, the pathogen test has posed substantial challenges in terms of biosecurity issues, times, costs and environmental impacts (Pham et al. 2020; Nguyen 2014). Due to these limitations, genomic selection has emerged as an alternative option to increase resistance of striped catfish to *E. ictaluri*, one of the most severe diseases that has caused significant economic loss for the sector world-wide.

Therefore, the principal of this study was to assess genomic prediction accuracies of machine and deep learning methods for two main disease resistance traits (i.e., survival status and survival time) and to make comparisons with four common methods (PBLUP, GBLUP, ssGBLUP and BayesR). Additionally, we explored if imputation of missing genotypes, multiple traits analyses and significant genome-wide markers can improve the prediction accuracies for the disease resistant traits. Our results open new opportunities for genome-based selection to increase the animal resistance to *E. ictaluri* in striped catfish as well as other aquaculture species infected by this highly infectious pathogen.

## 2. Materials and Methods

### Ethical statement

All the methods and experimental protocols of this study were performed in accordance with guidelines and regulations approved by the animal ethics committee of the University of the Sunshine Coast, Australia (approval number ANE1826).

#### 2.1. Fish and challenge test

The animal samples used in this study originated from a selective breeding program for improved disease resistance of striped catfish to *E. ictaluri* (Vu et al. 2019a). In 2020, the first generation was produced based on a nested mating design with a ratio of one male to two females. A total of 166 families (32 full- and 134 half-sib families) were successfully produced following the breeding protocol as detailed in Van Sang et al. (2012) and Vu et al. (2019b). Fry of each family were kept in separate fiber glass tanks up to 3 weeks before they were transferred to stock in net hapa installed in earthen pond. When the fingerlings reached an average body weight of 15 – 20 g, a random sample of 100 fish per family was individually identified using PIT (Passive Integrated Transponder) tag. After tagging, a half of each family was sent to ponds for performance testing and another half was used in pathogen challenge tests for *E. ictaluri* resistance.

The challenge test involved a total of 5328 individuals from 166 families (averaging 32 individuals per family). The experimental fish were initially acclimatized in cement tanks for about two weeks. Then the same number of fish from each family was randomly allocated to six cement tanks (10 m^3^) for the challenge test using cohabitant method (Vu et al. 2019a). The cohabitant fish (16.7 ± 6.1 g) were firstly inoculated with the bacteria *E. ictaluri* pathogen (10^6^ CFU/0.2 ml per fish). Two days after the injection, they were released into the cement tanks to rear with the experimental fish with a ratio of 1 to 3 (or roughly 30% cohabitant fish in each tank). The experiment was conducted over a period of 23 days. During this period, the feeding rate was reduced to 1.5% of the total biomass in tank. Mortality was highest in day 5 and dead fish were sampled for laboratory PCR test to verify that their death symptoms (white spots in spleen, liver and kidney) were due to *E. ictaluri* pathogen. At the conclusion of the experiment, all alive fish were biosecure-buried, following the regulations of the national veterinary authority (Department of Animal Health, Vietnam).

#### 2.2. Phenotype data

During the challenge test, death fish were collected every 3 hours and their clinical symptoms were also recorded. The data were used to calculate two measures of *E. ictaluri* resistance, i.e., survival status and survival time. Survival status was expressed as a binary trait in which dead fish were designated as zero (0) and alive animals at the end of the challenge test were assigned a number 1. Furthermore, survival time was defined as the continuous trait from the start of test until the animal death in day. Both survival status and survival time were analyzed using linear mixed model by ASReml 4.1 (Gilmour et al. 2014) to estimate breeding values (EBVs) for all individuals (5,328) and families (166) in the pedigree, with common environmental effect – *c^2^* (i.e., accounting for differences due to separate rearing of families until tagging) fitted in the model. Based on the EBVs ranking, 20 highest resistance families and 20 lowest resistance families were chosen from 166 families. Only families with the number of offspring greater than 9 were selected to ensure that the EBVs were estimated with a high level of reliability. Next, we randomly collected fin tissue samples of 12 – 15 fish per family per disease resistance group for genome sequencing using Diversity Arrays Technology (DArTseq™).

#### 2.3. Genotyping and quality control

A total of 564 DNA samples (from 564 individual fish) were sent to a commercial service provider in Canberra, Australia for genotyping by sequencing using DArTseq™ technology. The DArTseq™ was based on the genome complexity reduction method in combination with high throughput next generation sequencing using Illumina platform. The sequencing protocol including choice of restricted enzymes and library preparation was optimized for striped catfish (Pangasianodon DarTseq 1.0) using 96 independent DNA samples of striped catfish in our earlier study (Vu et al. 2020). Briefly, the PstI-SphI method (Kilian et al. 2012) was used and the compatible adaptors were designed to include Illumina flow-cell attachment sequence, sequencing primer sequence and capturing variant length of barcode regions (Elshire et al. 2011). Then, only mixed fragments (PstI-SphI) were amplified in 30 rounds of PCR, followed by sequencing on Illumina Hiseq2500 (77 cycles per single read). Next, sequences generated in each plate with 96-well microliter were processed by proprietary DArT analysis protocol. The genotype calling were partially described in our previous studies (Nguyen et al. 2020; Nguyen et al. 2018). The average variant call-rate was 99% and the sample calling rate was 92%. With this quality control, four out of 564 samples were discarded, and 560 samples (490 offspring and 70 parents) and 14.154 SNPs remained for further quality control by dartR package (Gruber et al. 2018). In this data, average read depth in each locus ranged 2,800 to 122,106 sequences, reads count from 1,410 to 81,949 sequences. The quality control of the genotype data in the dartR package (Gruber et al. 2018) was filtered for loci call-rate (<0.05, 4665 SNPs removed) and individual call-rate (<0.9, 2 individuals removed), monomorphic SNPs (0 SNP removed), minor allele frequency (<0.05, 2830 SNPs removed) and significant SNPs departure from Hardy-Weinberg Equilibrium (<0.05, 0 SNP removed). The retaining 6659 SNPs were blasted to the striped catfish genome database GENO_Phyp_1.0 (Kim et al. 2018) which matched 6470 SNPs to the genome chromosome information. The genotype data of 6470 SNPs and the phenotypes of 488 individuals were used for subsequent analyses.

#### 2.4. Genotype imputation

The missing genotypes were about 10.0% in this study. They were imputed using AlphaFamImpute (Whalen et al. 2020). The imputation was based on offspring-parent (i.e., pedigree) information. Specifically, this method used parental genotypes to impute the missing values of offspring genotypes and then, the offspring genotypes were used to fulfill the missing values of their parents (Whalen et al. 2020). The pedigree used in our analysis was traced back to the base population, including 4 generations.

#### 2.5. Estimation of heritability using genotype data

Variances components of the two studied traits (survival status and survival time) were estimated using linear mixed model under the Best Linear Unbiased Prediction (BLUP) framework in AIREMLf90 sub-program of the BLUPF90 family package (Misztal et al. 2002). AIREMLf90 uses Average Information-Restricted Maximum Likelihood method that requires less computational resources and higher accuracy of variance estimates (Masuda et al. 2014). The models included the fixed effects of spawning batch (4 levels, 1 – 2 weeks interval between successive spawning batches) and pathogen challenge test tanks (2 levels) and the random effect of additive genetics of the individual fish in the pedigree. Age from birth to tagging (124 – 167 days) was also fitted as a linear covariate. The SNP heritability (*h*^2^) was calculated as a ratio of the additive genetic variance 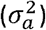 to total phenotypic variance 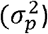, where 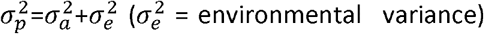. Based on the logarithmic likelihood ratio test (LRT), the common environmental effect (*c*^2^) was not significant (P > 0.05) and hence, this effect was omitted from the statistical models used to estimate genetic variances and prediction accuracy.

#### 2.6. Statistical methods

Accuracies of genomic prediction for the disease resistance traits (survival status and survival time) were estimated using BLUP-family methods (PBLUP, GBLUP, ssBLUP), BayesR, machine learning (i.e., KAML) and deep learning (i.e., DeepGP).

##### 2.6.1. PBLUP, GBLUP and ssGBLUP (using BLUPF90 program)

The mixed model used in PBLUP, GBLUP and ssGBLUP is written in a matrix notation as follows:

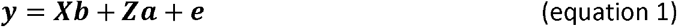

Where ***y*** is the vector of phenotypic values (survival status or survival time), ***b*** is the vector of fixed effects (i.e., spawning batches and experimental tanks and a linear covariate of age from birth to tagging), ***a*** is the vector of the random term (i.e., the additive genetic effect of individual fish in the pedigree). ***X*** and ***Z*** are the incidence matrices related to the fixed and random effects. The letter **e** refers to residual variance or error of the estimates corresponded to each phenotypic value.

The main difference among BLUP-family methods (PBLUP, GBLUP and ssGBLUP) relates to relationship matrices (**A, G** or **H**) used to solve the mixed model equation 1 above.

###### PBLUP

In PBLUP (Henderson 1985), ***a*** is the additive genetic effect of each individual fish with its corresponding matrix **Z** following a normal distribution 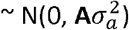, with **A** is the numerator relationship matrix calculated from the pedigree records and 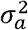 is the additive genetic variance.

###### GBLUP

In GBLUP (Meuwissen et al. 2001), ***a*** is the random additive genetic effect underlying polygenic effects of SNPs with a normal distribution 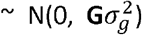 where genomic relationship matrix **G** was generated by VanRaden method (VanRaden 2008) from 6,470 SNPs and genomic variance 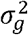. The GBLUP method assumes that each SNP contributes little to phenotypic variance of the traits studied.

###### ssGBLUP

In ssGBLUP (Misztal et al. 2009; Lourenco et al. 2020), **A** and **G** matrices are blended to produce realized matrix **H** (Aguilar et al. 2010). The inverse of H matrix is expressed as below:

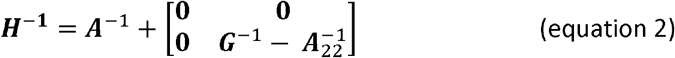

Where **A**^−1^ and **G**^−1^ are the inverse of matrices **A** and **G** as described above. **A**^−1^_22_ is the inverse of matrix of genotyped individuals only.

The three BLUP methods (PBLUP, GBLUP and ssGBLUP) were implemented in BLUPF90 package (Misztal et al. 2002). For survival status, we used generalized (threshold) mixed model in THRGIBBF1f90 (Tsuruta and Misztal 2006) known as threshold Gibb Sampling method, whereas survival time was analyzed using linear mixed models in AIREMLf90 (Masuda et al. 2014). In the Gibbs sampling, we used 200,000 iterations with 20,000 iterations as burn-in for univariate analysis. The convergence of the parameters estimates was checked using POSTGIBBSf90 program.

##### 2.6.2. Non-linear Bayesian method (BayesR)

BayesR assumes that SNP effects *β_i_* follow a four-component normal mixture and the effect of SNP *i* is assumed to be distributed as:

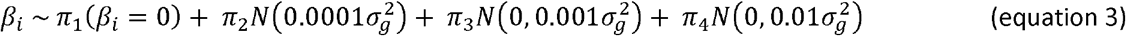

Where 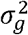 as defined above, 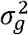 represents the total additive genetic variance (i.e., the cumulative variance of all SNP effects) and *π* = (*π*_1_, *π*_2_, *π*_3_, *π*_4_) the mixing proportions such that 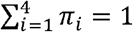. The mixing proportions *π* are assumed to follow a Dirichlet prior, *π ~ Dirichlet* (*α* + *β*), with *α* representing a vector of pseudo counts and *β* the cardinality of each component. In this work, we used a flat Dirchlet distribution, with *α* = (1,1,1,1) for the prior. As suggested by Moser et al. (2015), 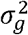 is assumed to be a random variable following an Inverse—χ2 distribution.

The analysis of BayesR consisted of three mains steps. First, numeric coding genotypes of 560 animals (including 490 offspring and 70 parents) obtained from DArT sequencing were converted to ATGC format (i.e., biallelic variant). Second, the trimmed sequences of each variant were blasted to nucleotide non-redundant striped catfish database using Blast2GO (Conesa et al. 2005), named GENO_Phyp_1.0 (Kim et al. 2018). This analysis matched 13,766 SNPs to 30 autosomal chromosomes. The remaining SNPs (388 SNPs) were randomly assigned to 30 chromosomes (12 – 13 SNPs/chromosome). Third, PLINK v1.9 (Purcell et al. 2007) that utilizes genotype and mapping files was used to generate a full dataset for BayesR (https://github.com/syntheke/BayesR). The default running cycles of BayesR included 50,000 steps with 20,000 steps discarded as burn-in. We used predicted phenotypes (computed in AIREMLf90 using equation 1), as this package does not allow fitting covariates in the model.

##### 2.6.3. Machine learning and deep learning methods

###### KAML (Kinship Adjusted Multiple Loci)

KAML is a machine learning-based method that incorporates cross-validation, multiple regression, grid search and bisection algorithm to improve genomic prediction accuracy for complex quantitative traits (Yin et al. 2020). KAML extends linear mixed model (equation 1) by incorporating quantitative trait nucleotides - QTNs (e.g., traits controlled by genes with large and moderate effects) as covariates and a SNP-weighted trait-specific kinship matrix as the (co)-variance assumption corresponding to the random effect. The selections of QTNs and SNP weights are optimized by machine learning. In a matrix notation, equation 1 becomes:

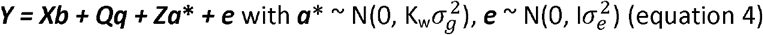

Where 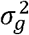 and 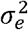 are as described above; ***Q*** is covariates matrix of QTNs detected throughout multiple regression analyses; ***q*** is a vector of fixed effect of each QTN corresponding to ***Q*** and K_w_ is SNP-weighted Kinship that is optimized by grid search and bisection procedure. The implementation of KAML included two main steps, i.e., define ***Q*** and K_w_ and then predicting genomic breeding values of the validation set. In the training step, KAML optimized parameters for determining QTNs and SNPs weights which used all testing subset data provided throughout cross-validation. Next, in the prediction step, the optimized parameters were used to predict genomic breeding values of each individual in validation set. A detailed description of the step-wise procedures and the underlying algorithms of KAML is given in Yin et al. (2020). In our analyses, we used PLINK (Purcell et al. 2007) to convert the genotype data into the right format to be analyzed in KAML (https://github.com/YinLiLin/KAML) in R environment (R Core Team 2015). KAML utilized adjusted phenotypes (i.e., phenotypic y-hats) for testing sets and masked (i.e., ‘NA’ values) phenotypes of validation sets. To define the best parameters (i.e., pseudo QTNs, the optimal SNPs weights, the optimal Log value and the optimized kinship matrix) for mixed models of each testing set, we used five-fold cross-validation with 100 replications (to obtain good model estimates).

###### Deep learning – Multilayer Perceptron (DL-MLP)

For deep learning analysis, we used DeepGP package (https://github.com/lauzingaretti/deepGP/). A detailed description of the theoretical framework is given in Zingaretti et al. (2020) and Pérez-Enciso and Zingaretti (2019). In our analyses, we used two different algorithms of deep learning: Multilayer perceptron and convolutional neural network.

Multilayer perceptron is fully connected networks consisting of an input layer, one or several hidden layers, and an output layer. In the context of genomic prediction, input layer receives SNPs data (i.e., 6,470 SNPs). Each of SNP information in the input layer is transferred its value to the first neuron of the first hidden layer by the Z function:

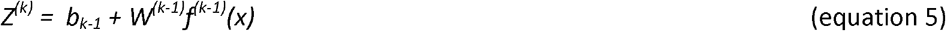

Where *W* is weighted SNP information, *f*(*x*) is non-linear function and *b* is bias (i.e., constant). In the case of genomic prediction, the input layer is SNPs genotype (i.e., input variables = number of SNPs) of individual fully connected to the first hidden layers. As a result, the value in the first neuron of the first hidden layer is the sum effect of Z function over 6,470 SNPs 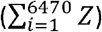. Likewise, information in the first hidden layer is the input for transferring its value to the second hidden layers and finally to the output layers. The significance step in obtaining best DL-MLP model is to define MLP structure network (i.e, number of hidden layer and neuron of each layer) and hyperparameters which related to equation 5. These parameters and their values in our study include learning rate (--lr 0.025 0.01), type of drop out (--dr 0 0.01), type of regularization (--reg 0.0001), activation function (--act linear tanh), number of hidden layer (--h 1 5), optimizer function (--optimizer Adam) or number of epochs (--epochs 50). These scripts provided in DeepGP (https://github.com/lauzingaretti/DeepGP) were used to obtain hyperparameters in our DL-MLP analyses.

###### Deep learning – Convolutional Neural Network (DL-CNN)

Convolutional neural network comprises of 3 types of layers: (1) convolutional layers, (2) fully connected layers and (3) input layers. In the context of genomic prediction, first convolutional layer receives data of 6,470 SNPs and write it to the first layer using kernel (i.e., a definite number of SNPs is grouped or combined). Then, the pooling step is responsible to reduce spatial size of SNPs data of the convolutional layer into the second convolutional layer. There are two types of pooling such as max pooling and average pooling corresponding to maximum values and mean values of SNPs portion covered by the kernel. The process was repeated to make update value of the last convolutional layer. At the last convolutional layer, data were flattened (i.e., convert data of pooled feature of the last convolutional layer into single column feeding to fully connected layers). The fully connected layer works similarly the MLP. The hyperparameters used in our study are size of kernel (--ks 3), learning rate (--lr 0.025 0.01), dropout in the first layer (-dr1 0 0.01), dropout in the hidden layer (--dr2 0 0.1 0.2), number of pooling size (--ps 1), number of convolutional layers (--ncov 1 3), number of strides (--ns 1), number of convolutional operations (--nfilters 8). These hyperparameters are fine-tuned under DeepGP with grid search provided in DeepGP package (Zingaretti et al. 2020). As DeepGP cannot handle missing values, we used imputed genotype for all analyses.

##### 2.6.4. Analysis of highly significant SNPs and multi-trait mixed model

###### ssGWAS to select significant SNPs for prediction

Single step genome-wide association analysis (ssGWAS) was accomplished in BLUPF90 suite using each subset phenotype data for both disease resistance traits. We used preGSf90 and postGSf90 together with BLUPF90 program to compute SNP effect and its P-values. SNPs panels were selected based on the P-value < 0.0001 (the number of the selected SNPs sets were presented in Supplementary Table S2). These SNP panels were used to test the predictive performance of six genomic prediction methods used in our study.

###### Multi-trait mixed model

Multi-trait analyses are only available in BLUPF90. In this study, two studied traits were jointly analyzed using PBLUP, GBLUP and ssGBLUP models in THRGIBBS1F90 (Tsuruta and Misztal 2006). This involves using Gibbs Sampling with MCMC updates including 1,000,000 iterations with 100,000 burn-in steps.

#### 2.7. Five-fold cross validation

The predictive performances of all seven methods (PBLUP, GBLUP, ssGBLUP, BayesR, ML-KAML, DL-MLP and DL-CNN) used in this study were evaluated using repeated 5-fold cross validation. The procedure involved a random partitioning of the full data into 5 subsets (each subset has 97 – 98 individuals including representatives of all the families). The phenotypes of one subset were masked and GEBV/EBV (Genomic/Estimated Breeding Value) of one set were predicted using the information from the other four subsets. This means one phenotype was used once in the validation process of one repetition. Each analysis was repeated 5 times, giving 25 validation sets in total and the correlation coefficient was averaged over 25 values. To compute evaluation metric in each validation set, the GEBV/EBVs of the predicted phenotypes were correlated to the actual phenotype. We chose Pearson correlation coefficient (Benesty et al. 2009) as an evaluation metric to consistently compare the statistical models used. Moreover, we computed Mathew correlation coefficient, MCC (Matthews 1975) for binary trait (survival status) as MCC is giving a different look (indicate which predicted value is assigned to 0 or 1 case) for binary classification evaluation (Chicco and Jurman 2020). Specifically, we used *PresenceAbsence* R package (Freeman and Moisen 2008) to produce 4×4 confusion matrix which assigned GEBV/EBV values against the actual phenotypes for computing MCC (Supplementary file Table S1 for additional information and reference). To obtain the consistency and comparable results among tested software/packages, we used the same subsets of testing/validation data in BLUPF90 family program, BayesR, ML-KAML, DL-MLP and DL-CNN.

The genomic prediction accuracy for survival status and survival time was calculated as 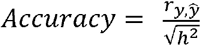; where 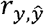 is the correlation between the predicted breeding values 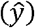 and actual phenotypes(*y*).

To evaluate reliability (or potential biases) of the statistical methods, we computed RMSE (Root Mean Square Error) and R^2^ (coefficient of determination) based on 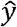 and *y* values. RMSE is the standard deviation of residuals, also known as prediction error, while coefficient of determination is an indicator of the goodness of fit of a statistical model. RMSE and R^2^ were calculated as below:

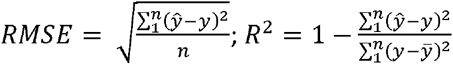

Where 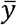 is the average value of *n* actual phenotypes in a testing set.

##### Data availability

Data are available via Figshare portal provided by the journal.

## 3. Results

### 3.1. Data, variance component and heritability

The average survival rate and survival time after the pathogen challenge test were 32.2% and 163.9 hours, respectively (Table 1). When exposed to the *E. ictaluri* pathogen, mortality occurred after 32.5 hours and the highest death rate was observed after 97.3 hours (Figure 1). There were significant differences (P < 0.0001, Figure 1) in both survival rate (60.0% vs. 4.5%) and survival time (207.8 vs. 120.0 hours) between 20 highest and 20 lowest resistant families (Table 1). The variations in the disease resistant traits among families were also significant (Supplementary Figures S1 & S2).

**Table 1.** Number of animals (n) and phenotypes in the disease resistant and susceptible groups used for genotyping.

| Groups | 20 Lowest survival family |  | 20 Highest survival families |  | All 40 families |  |
| --- | --- | --- | --- | --- | --- | --- |
| Trait | Individuals | Mean ± SD | Ind/family | Mean ± SD | n | Mean ± SD |
| Survival status (%) | 245 | 4.5 ± 20.8 | 245 | 60.0 ± 49.1 | 490 | 32.2 ± 46.8 |
| Survival time (hour) | 245 | 120.0 ± 52.8 | 245 | 207.8 ± 103.9 | 490 | 163.9 ± 93.4 |

**Figure 1.**
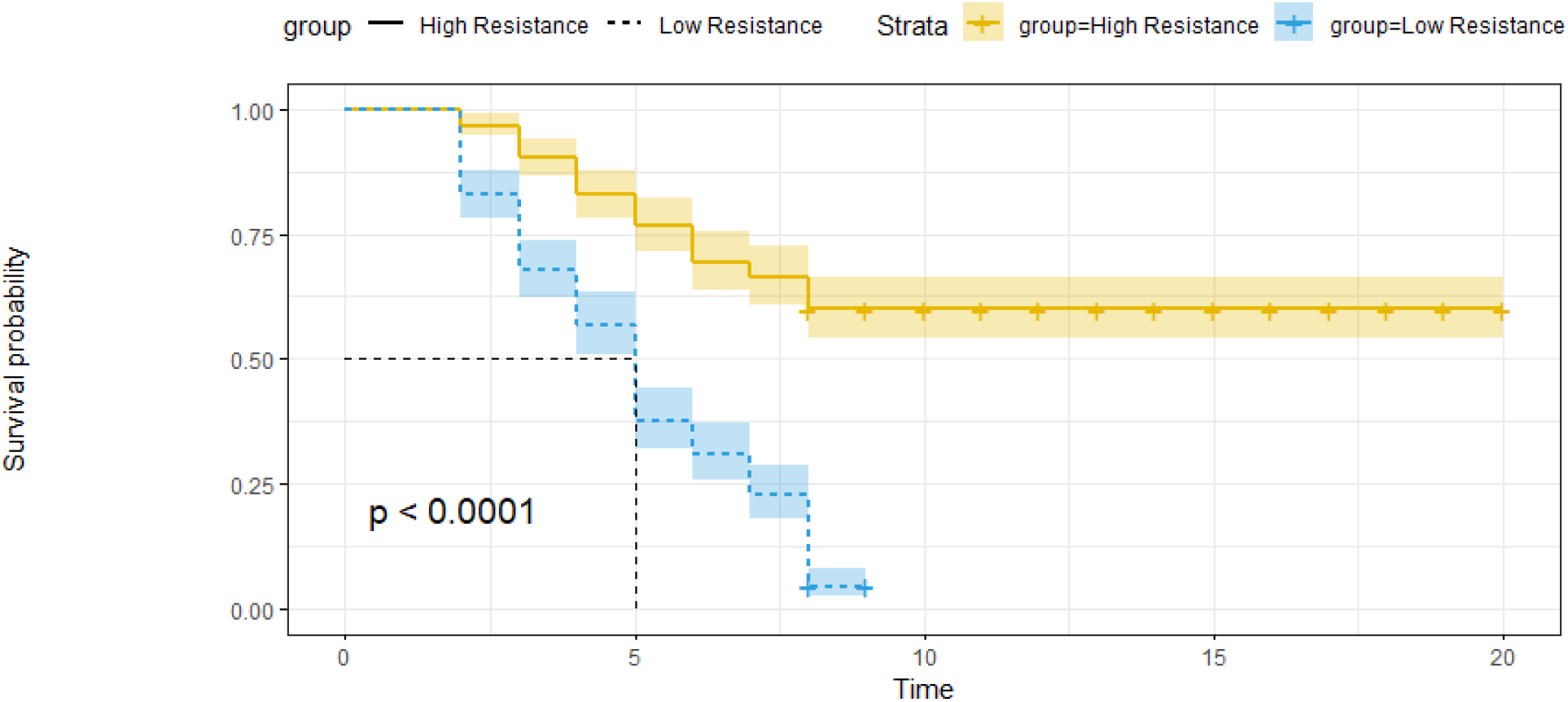
Survival trend in day post challenge of high and low resistance groups in genotyped individuals of striped catfish.

The heritabilities estimated for survival status and survival time using traditional PBLUP (0.65 – 0.71) were greater than those obtained from (ss)GBLUP (0.44 – 0.46) but they were lower than those estimated by BayesR (0.96 – 0.97) (Table 2). Across statistical models (methods) used, the estimates of heritability for survival status were slightly higher than those obtained for survival time (Table 2).

**Table 2.** Variance components and heritability for survival status and survival time using AIREMLf90 and THRGIBBS1f90 sub-program in BLUPF90 and BayesR

| Trait | Models | Additive genetic variance | Phenotypic variance | Heritability |
| --- | --- | --- | --- | --- |
| Survival status | PBLUP | 0.147 | 0.207 | 0.71 ± 0.14 |
|  | GBLUP | 0.086 | 0.188 | 0.46 ± 0.09 |
|  | ssGBLUP | 0.088 | 0.190 | 0.46 ± 0.09 |
|  | BayesR | 0.043 | 0.044 | 0.96 ± 0.03 |
| Survival time | PBLUP | 9.579 | 14.694 | 0.65 ± 0.14 |
|  | GBLUP | 5.900 | 13.303 | 0.44 ± 0.09 |
|  | ssGBLUP | 5.908 | 13.442 | 0.44 ± 0.09 |
|  | BayesR | 2.747 | 2.818 | 0.97 ± 0.02 |

Note that estimate of genetic variance was not provided by ML-KAML, DL-MLP and DL-CNN methods.

### 3.2. Accuracy of genomic prediction using original genotype (un-imputed) data

Figure 2 presents the accuracies of genomic breeding values for survival status and survival time using Machine learning (ML-KAML) and Deep Learning (DL-MLP and DL-CNN) together with four other methods (PBLUP, GBLUP, ssGBLUP and BayesR). When the original genotype data (i.e., un-imputed data) were used, the prediction accuracies from ML-KAML (0.67) was slightly higher than PBLUP (0.66) and they both were higher than those calculated from GBLUP (0.55) and ssGBLUP (0.55) for survival status. ML-KAML also outperformed BLUP methods for survival time (0.69 vs 0.59-0.65) (also see Supplementary Table S1). Additionally, the difference between ML-KAML and BayesR in the prediction accuracy was small (0.67 for survival status and 0.69 vs. 0.70 for survival time). For both traits, ssGBLUP was slightly better GBLUP. Interestingly, both GBLUP and ssGBLUP did not improve the prediction accuracies for both survival status and survival time as compared with conventional PBLUP method.

**Figure 2.**
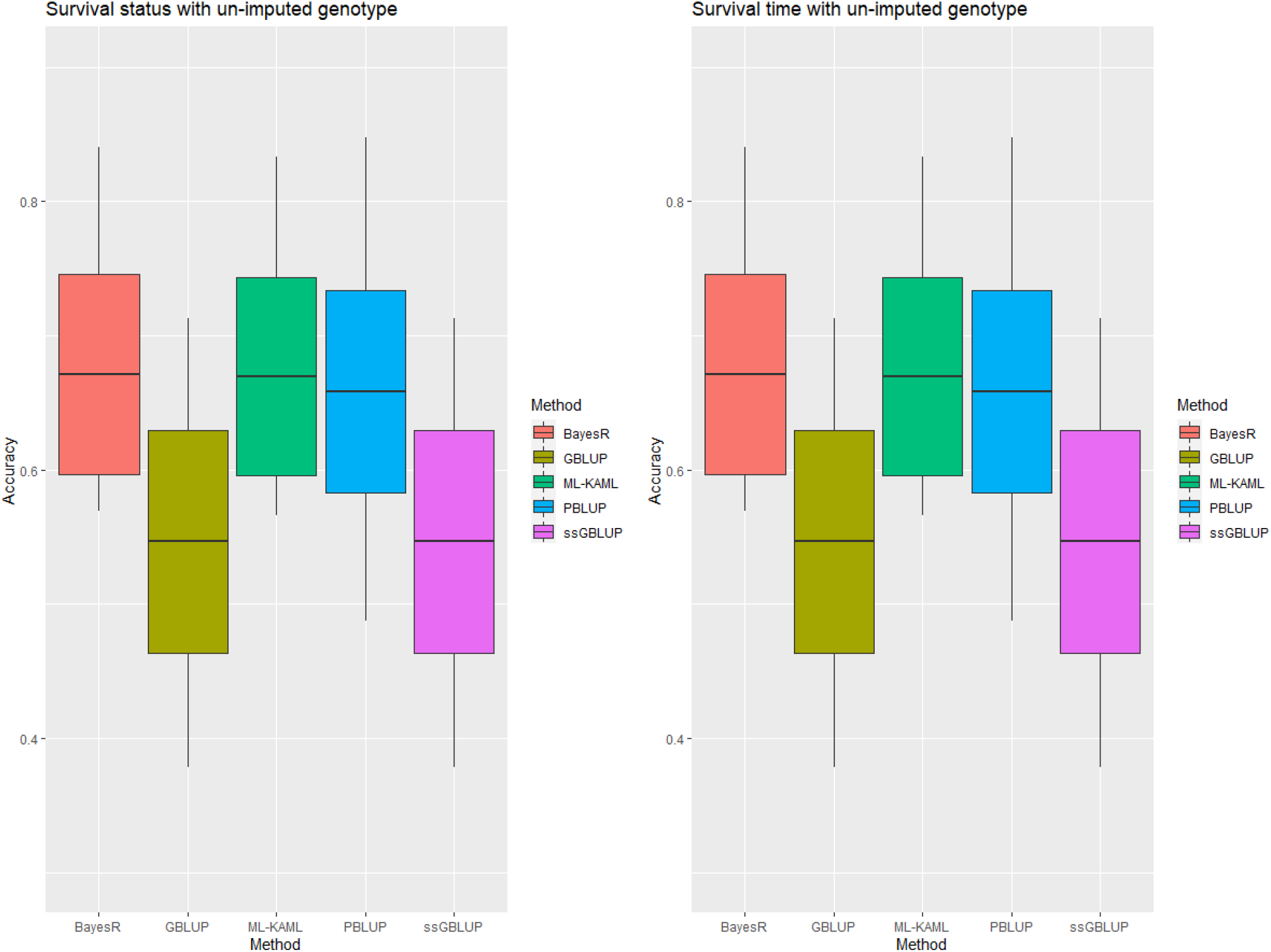
Accuracy of survival status and survival time traits using original genotype. Middle line of the box is mean accuracy; top and bottom lines of the box is accuracy ± one standard deviation. End points of vertical line represent min and max values. Note that PBLUP uses phenotype and pedigree information only.

### 3.3. Accuracy of genomic prediction using imputed genotype data

Imputation of missing genotypes increased the accuracies of genomic prediction for both survival status and survival time across the seven methods used (Figure 3). The improvements for survival status were from 12.0% (ML-KAML) to 19.2% (GBLUP). For survival time, the increase in the prediction accuracy when the imputed genotypes were used as compared with un-imputed data was largest for GBLUP (10.0%), followed by ML-KAML (8.1%), ssGBLUP (7.1%) and BayesR (5.3%). Deep learning (DL) using multilayer perceptron (DL-MLP) and convolutional neural network (DL-CNN) had lower prediction accuracies for both traits (0.67 – 0.73) than ML-KAML (0.75) or BayesR (0.75-0.76) methods but they (DL-MLP and DL-CNN) had higher prediction accuracies than BLUP methods (0.55 – 0.66) (Supplementary Table S1). ML-KAML and BayesR methods also had least variation (Min – Max values of accuracy in the testing set) compared to other methods (Figure 3).

**Figure 3.**
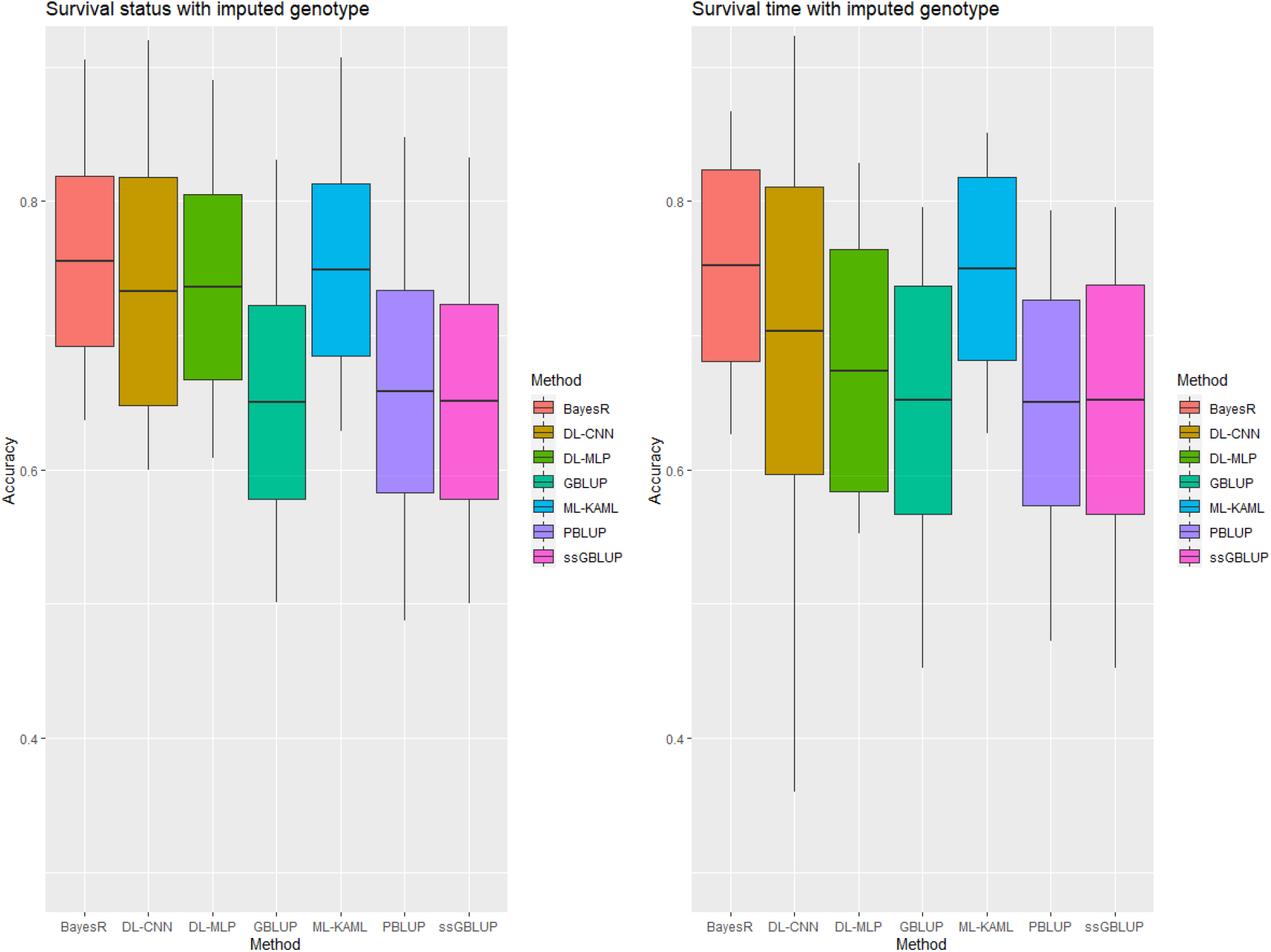
Accuracy of survival status and survival time traits using imputed genotype. Middle line of the box is mean accuracy; top and bottom lines of the box is accuracy ± one standard deviation. End points of vertical line represent min and max values. Note that PBLUP uses phenotype and pedigree information only.

### 3.4. Univariate vs. multi-variate analysis

Our multivariate analyses used PBLUP, GBLUP and ssGBLUP, and the prediction accuracies for both survival status and survival time are shown in Table 3. There were slight decreases observed in the prediction accuracy when the three BLUP methods were used to estimate genomic breeding values (GEBV) in multivariate analyses relative to univariate model. The reduction in the prediction accuracies was only 2.9 to 5.6% for survival time and 0.3 to 2.5% for survival status.

**Table 3.** Genomic prediction accuracy for survival status and survival time using univariate and bivariate models with un-imputed 6470 SNPs

| Trait | Method | Univariate |  |  | Bivariate |  |  | Increasement (%) |
| --- | --- | --- | --- | --- | --- | --- | --- | --- |
|  |  | Accuracy±sd | RMSE | R<sup>2</sup> | Accuracy±sd | RMSE | R<sup>2</sup> |  |
| Survival status | PBLUP | 0.66 ± 0.08 | 0.39 | 0.30 | 0.66 ± 0.07 | 0.39 | 0.30 | - 0.3 |
|  | GBLUP | 0.55 ± 0.08 | 0.41 | 0.21 | 0.53 ± 0.08 | 0.42 | 0.20 | - 2.5 |
|  | ssGBLUP | 0.55 ± 0.08 | 0.41 | 0.21 | 0.54 ± 0.08 | 0.41 | 0.21 | - 0.7 |
| Survival time | PBLUP | 0.65 ± 0.08 | 3.30 | 0.26 | 0.63 ± 0.09 | 3.33 | 0.25 | - 2.9 |
|  | GBLUP | 0.59 ± 0.09 | 3.41 | 0.22 | 0.57 ± 0.07 | 3.47 | 0.21 | - 3.5 |
|  | ssGBLUP | 0.61 ± 0.07 | 3.38 | 0.23 | 0.58 ± 0.07 | 3.47 | 0.21 | - 5.6 |

Note: bivariate analysis is not available in BayesR, ML-KAML, DI-MLP and DL-CNN. RMSE = Root Square Mean Error, R<sup>2</sup> = coefficient of determination, sd = standard deviation. Increasement (%) = (accuracy of bivariate – accuracy of univariate)/accuracy of univariate.

### 3.5. Genomic prediction in combination with GWAS

The prediction accuracies for the disease resistance traits that incorporated only significant SNPs identified from genome-wide association analysis (GWAS) are given in Tables 4 and 5. In all the genotype subsets used, the prediction accuracies were generally smaller to those of the full set of 6,470 SNPs. Using smaller SNP panels selected from ssGWAS with p<0.00001 in this study scarified accuracies by 0.3 to 15.6%. For survival status the prediction accuracy was reduced from 4.3 to 6.0% for ML-KAML and BLUP methods, but greater loss was observed for deep learning methods (11.2% for DL-CNN and 15.6% for DL-MLP). The same trend was observed for survival time but the reduction in the prediction accuracy was only 0.3 to 4.9%.

**Table 4.**
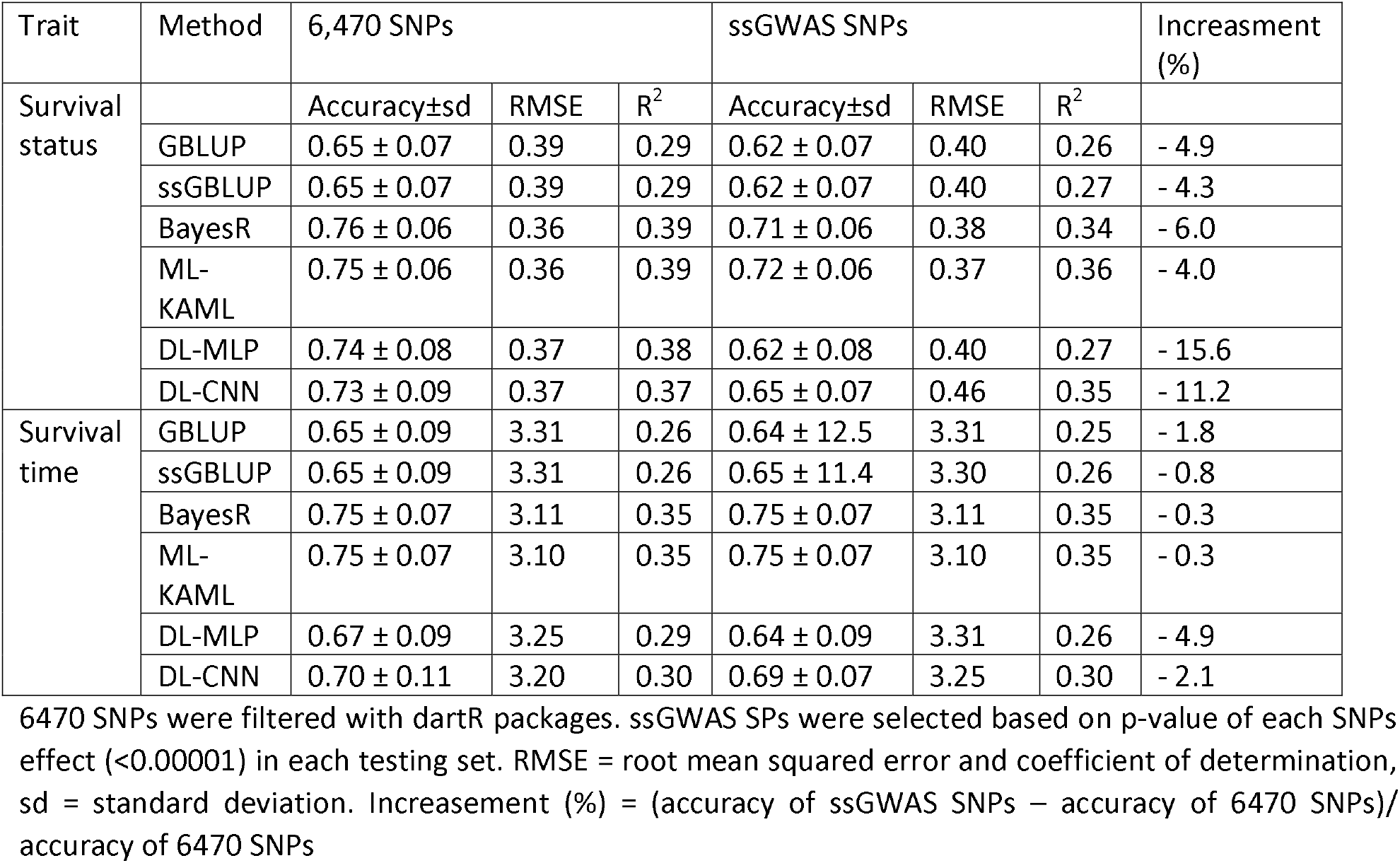
Accuracy of genomic prediction using imputed genotypes of 6470 vs ssGWAS SNPs.

**Table 5.**
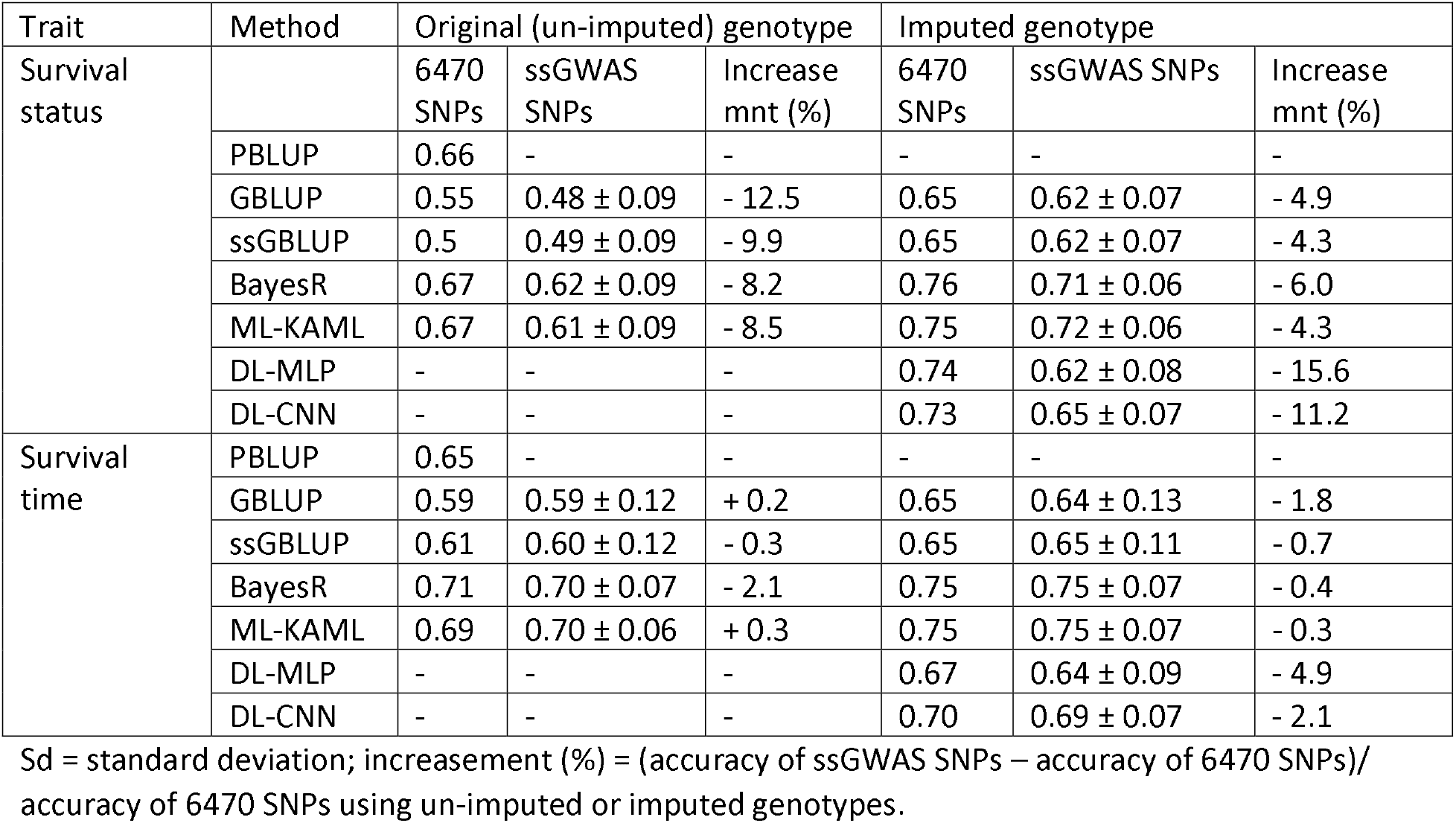
Accuracy (±sd) of genomic prediction for survival status and survival using highly significant SNPs (P < 0.0001)

| Trait | Method | Original (un-imputed) genotype |  |  | Imputed genotype |  |  |
| --- | --- | --- | --- | --- | --- | --- | --- |
| Survival status |  | 6470 SNPs | ssGWAS SNPs | Increase mnt (%) | 6470 SNPs | ssGWAS SNPs | Increase mnt (%) |
|  | PBLUP | 0.66 | - | - | - | - | - |
| | GBLUP | 0.55 | $0.48 \pm 0.09$ | - 12.5 | 0.65 | $0.62 \pm 0.07$ | - 4.9 |
| | ssGBLUP | 0.5 | $0.49 \pm 0.09$ | - 9.9 | 0.65 | $0.62 \pm 0.07$ | - 4.3 |
| | BayesR | 0.67 | $0.62 \pm 0.09$ | - 8.2 | 0.76 | $0.71 \pm 0.06$ | - 6.0 |
| | ML-KAML | 0.67 | $0.61 \pm 0.09$ | - 8.5 | 0.75 | $0.72 \pm 0.06$ | - 4.3 |
| | DL-MLP | - | - | - | 0.74 | $0.62 \pm 0.08$ | - 15.6 |
| | DL-CNN | - | - | - | 0.73 | $0.65 \pm 0.07$ | - 11.2 |
| Survival time | PBLUP | 0.65 | - | - | - | - | - |
| | GBLUP | 0.59 | $0.59 \pm 0.12$ | + 0.2 | 0.65 | $0.64 \pm 0.13$ | - 1.8 |
| | ssGBLUP | 0.61 | $0.60 \pm 0.12$ | - 0.3 | 0.65 | $0.65 \pm 0.11$ | - 0.7 |
| | BayesR | 0.71 | $0.70 \pm 0.07$ | - 2.1 | 0.75 | $0.75 \pm 0.07$ | - 0.4 |
| | ML-KAML | 0.69 | $0.70 \pm 0.06$ | + 0.3 | 0.75 | $0.75 \pm 0.07$ | - 0.3 |
| | DL-MLP | - | - | - | 0.67 | $0.64 \pm 0.09$ | - 4.9 |
| | DL-CNN | - | - | - | 0.70 | $0.69 \pm 0.07$ | - 2.1 |

Sd = standard deviation; increasement (%) = (accuracy of ssGWAS SNPs – accuracy of 6470 SNPs)/accuracy of 6470 SNPs using un-imputed or imputed genotypes.

### 3.6. Reliability (or potential biases) of the statistical methods

The root mean square error (RMSE) and coefficient of determination (R^2^) were computed for each method and they are shown in Tables 3 and 4. In both traits (survival status and survival time) and in all analyses (univariate and multi-trait or full genotype vs subsets), the machine learning (i.e., ML-KAML) gave the least possible biases in the breeding values prediction, as evidenced by the low RMSE (0.36 – 0.37 for survival status and 3.10 for survival time) and high R^2^ (0.35 – 0.39). BayesR also displayed similar predictive reliability to ML-KAML (RMSE = 0.36 – 0.37 for survival status and 0.311 for survival time and R^2^ = 0.34 – 0.39). Among BLUP family methods, there were no large differences in the reliability of the genomic breeding value estimation between GBLUP and ssGBLUP (for instance with univariate model using un-imputed genotype, RMSE = 0.41 and R^2^ = 0.20 – 0.21). However, they were likely less reliable than PBLUP (RMSE = 0.39 and R^2^ = 0.30). The utilization of the imputed genotypes in GBLUP and ssGBLUP methods reduced the prediction errors for both survival status and survival time (RMSE = 0.39 and 3.31, and R^2^ = 0.29 and 0.26, respectively).

## 4. Discussion

Development of disease resistance lines is essential for the aquaculture sector due to the combined effects of aquaculture intensifications and environmental changes. Our study demonstrated that genomic selection can be practiced improving resistance to *Edwardsiella ictaluri* disease in striped catfish populations, using a range of different statistical methods especially machine and deep learning, and BayesR. Salient findings from our study are discussed as follows.

First, the genomic prediction accuracies for the disease resistance traits (survival status and survival time) were substantially improved when machine (i.e., ML-KAML) and deep learning (DL-MLP, DL-CNN) methods were used relative to BLUP-family methods (PBLUP, GBLUP, ssGBLUP). In addition, our evaluation of the prediction biases or prediction errors based on RMSE and R^2^ statistics indicated that the AI methods may be prone to the least errors in the estimated breeding values than BLUP family methods. Hence, genomic selection to improve *E. ictaluri* disease is expected to achieve greater genetic gain than the conventional selective breeding approach using pedigree and phenotype information. The genomic selection can also shorten the breeding cycle of striped catfish as expected to reduce at least a half of generation interval in dairy cattle (Hayes et al. 2009). Our results are consistent with those reported in farmed animal (Baker et al. 2020), livestock (Li et al. 2018; González-Recio and Forni 2011) and plants (Montesinos-López et al. 2019; Montesinos-López et al. 2018; Zingaretti et al. 2020). Despite the superiority of the machine learning methods (i.e., ML-KAML) to PBLUP, GBLUP and ssGBLUP, they had similar predictive power to BayesR for both traits in our study. Reports in farmed animals concluded that BayesR is the best amongst Bayesian and GBLUP methods (Erbe et al. 2012) but no published information is available in aquaculture species to compare with our study. To date, only some studies, one in shrimp (Palaiokostas 2021) that used extreme gradient boost method (one of the machine learning approaches) and another in seabream that used support vector machines and linear bagging classification (Bargelloni et al. 2021) and reported that these methods improved the prediction accuracy for survival traits by 1-4% and 20 – 70% relative to GBLUP and Bayesian methods. Recently, the evaluation of deep learning model (e.g., convolutional neural network) in Bay scallop indicated this method outperformed BayesB and random regression GBLUP methods (Zhu et al. 2021). Therefore, machine or deep learning should be used for genomic evaluation of disease resistance traits, at least in this population of striped catfish. The predictive powers of these methods using AI algorithms are expected to be greater when big data (e.g., hundreds of thousands or million animals sequenced) are analyzed for this population in the future.

Second, imputation of missing genotypes (using offspring and parent information available in the pedigree of striped catfish) increased the predictive ability of all the seven methods used, especially BayesR by 5.3% to approximately 19% (GBLUP and ssGBLUP) relative to when un-imputed data were used. Also note that the imputation of missing genotypes in our study differed from others that used software to compute missing genotypes without considering parental relationship. With the use of offspring-parent information, the imputation likely increased accuracy of G and H matrices used in GBLUP and single-step methods (i.e., ssGBLUP). Published information across species also showed that imputation from low to high density SNP panels or from commercial SNP arrays to whole genome sequence increased the genomic prediction accuracies for resistance to parasite (i.e. sea lice count) and body traits in Atlantic salmon (Tsai et al. 2017), and agronomic traits in maize (Crossa et al. 2013). However, some other studies also showed that imputed genotype did not improve prediction accuracies in livestock animal breeding (Heidaritabar et al. 2016; Chen et al. 2014; Van Binsbergen et al. 2015). Imputation has several benefits, firstly it can be made after sequencing with no extra cost. Second, through imputation, we can increase the number of animals genotyped, which in turn increase selection intensity, and this can lead to increases in genetic progress made in selected populations. Thus, imputation of low-density sequence can support our future incorporation of genomic selection for *Edwardsiella ictaluri* disease in this striped catfish population.

Third, we found that multivariate analysis did not improve the predictive ability for the two studied traits irrespective of the statistical methods. These results are expected due to the high and positive genetic correlations (*r_g_*) between the two traits (*r_g_* = 0.81). A bivariate analysis of harvest body weight and fillet yield in Nile tilapia also showed no improvement in the prediction accuracies for these traits (Joshi et al. 2020). Multi-trait analyses generally improve accuracy of genetic parameter estimates and animal breeding values for traits whose genetic architectures are different (e.g., low vs high heritable) or genetically antagonistic associated (i.e., unfavorably correlated). Examples of multivariate genomic prediction models involving multi-trait in cassava and wheat plants gave better prediction accuracy over univariate models (Okeke et al. 2017; Sun et al. 2017). To utilize the benefits of multi-trait analysis, we continue collecting other traits of economic importance in striped catfish, namely disease tolerance, disease resilience as well as immunological parameters to increase overall resistant capacity of the animals in this catfish population.

Fourth, we also suggest using the full set of SNPs to obtain a reasonable level of prediction accuracy for genomic selection to improve disease resistance traits in striped catfish. It has been well documented that the prediction accuracy increased with number of markers and sequencing depth or genome coverage. Due to characteristics of RAD-sequencing methods, there are still limited number of markers in our study as compared with those reported in farmed animals or plants (Robledo et al. 2018; Davey and Blaxter 2010). It is necessary to examine prediction ability of whole genome data when sequencing costs are reduced to an affordable level to sequence thousands (or hundreds of thousands) of animals in a near future. However, there are studies in farmed animals reporting that whole genome sequence data improved the predictive powers for complex traits by only 0.4 – 5.0% (Al Kalaldeh et al. 2019; VanRaden et al. 2017).

Finally, the utilization of highly significant SNPs from ssGWAS can give comparative prediction accuracies for the two traits studied, for both un-imputed and imputed genotype data, as compared to whole SNP panel. Using significant SNP panels selected from ssGWAS had higher accuracy in the estimated breeding values for disease resistance traits in *Litopeneaus vannamei* (Luo et al. 2021) or fish species (Jeong et al. 2020). On the one hand, these SNPs can be used to develop a small SNP panel to genotype a large number of animals in both the training and validation populations. On the other hand, the costs of genotyping animals for hundreds of SNPs may not be a lot cheaper than DArT sequencing as used in this study. The use of significant SNPs can be beneficial when their biological functions (pathways) are known. In species where reference genome or high quality genome assembly is available, the incorporation of SNPs with biological functions (i.e. multi-omics) improved the prediction accuracy, for instance, about 5 - 19% for fertility traits in dairy cattle (Abdollahi-Arpanahi et al. 2017; Nani et al. 2019), or 27.4 – 60.7% for traits in inbred lines of Drosophila (Ye et al. 2020). Unfortunately, a good genome assembly is currently not available for striped catfish (Kim et al. 2018), this area thus deserves future studies to enable the efficient utilization of genomic information in the genetic selection program for this economically important species in the aquaculture sector.

In summary, despite the high level of consistency in the prediction accuracies across the seven methods used, the estimates of heritability and prediction accuracies obtained for survival status and survival time are potentially biased upward due to the family structure of the current population that included selective genotypes (high and low families). However, this problem may have been alleviated because both pedigree and genotype data were included in mixed models (e.g., BLUP family methods) to estimate genomic breeding values (Gowane et al. 2019) or machine learning methods as demonstrated in this study.

## 5. Concluding remarks

The genomic prediction accuracies for disease resistance traits of striped catfish were moderate to high, suggesting possibilities for the application of genomic selection to improve resilience to *Edwardsiella ictaluri* in this population. Machine learning and deep learning methods outperformed BLUP-family in most of our analyses, regardless full or subset data or only significant SNPs used with imputed genotype. However, the prediction accuracies using machine learning and deep learning for both survival status and survival time were almost similar to those obtained from BayesR. There were no clear advantages of machine learning and deep learning to conventional methods (PBLUP, GBLUP and ssGBLUP) if original genotype was used for survival status but its superiority was evidenced for survival time. Therefore, either machine learning, deep learning or BayesR could be used for genomic evaluation of the disease resistance traits in this striped catfish population. Furthermore, breeding to improve resistance to *E. ictaluri* can use survival status or survival time as alternative selection criterion as there was no significant difference in the prediction accuracies across the seven different methods used in our study. However, when more data are accumulated in future generations, it is necessary to re-evaluate the prediction accuracies and potential biases before any practical implementation of genomic selection can be made to improve disease resistance of this stripped catfish population.

---

**Supplementary Figure S1.**
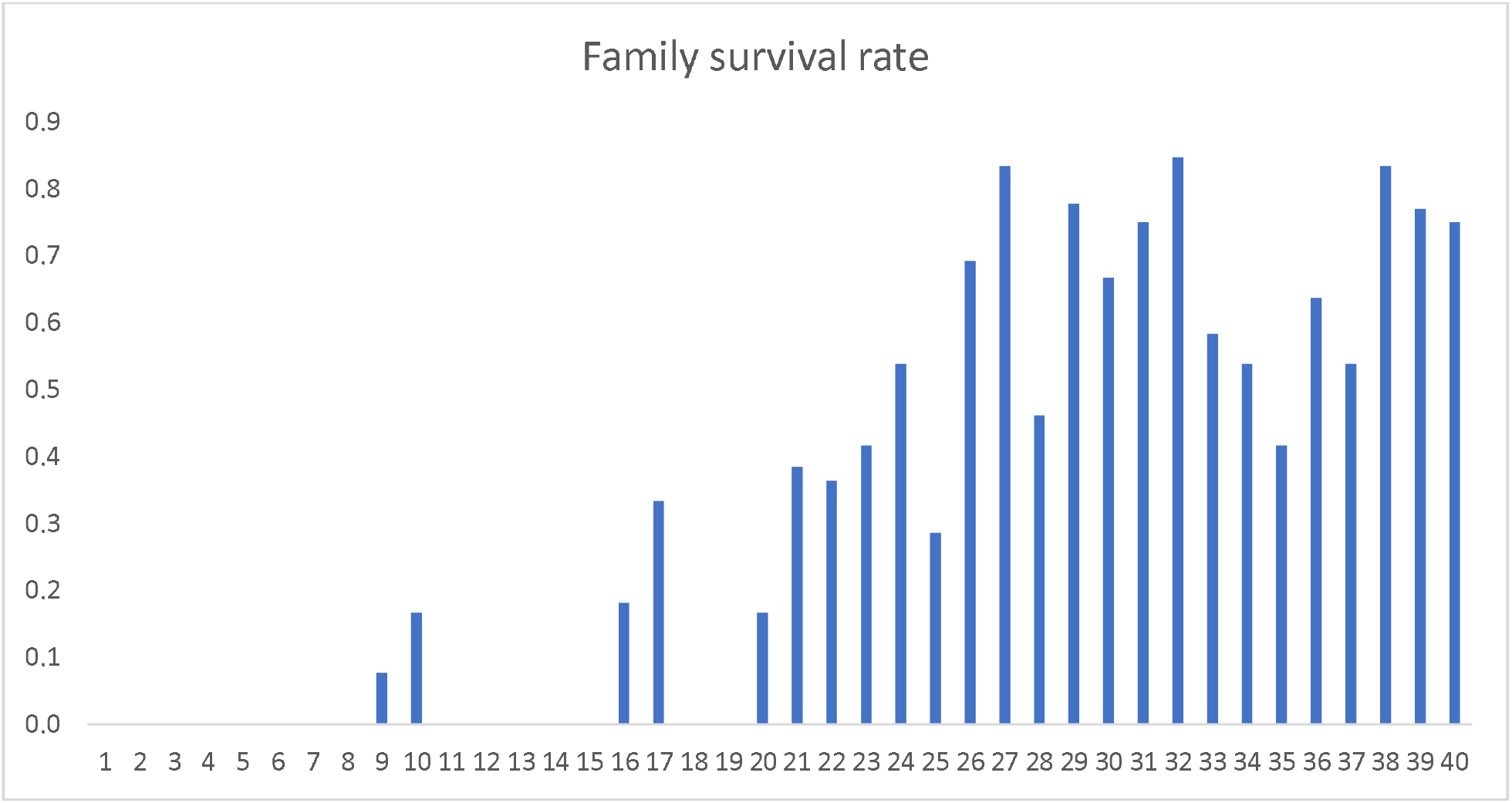
Mean family survival rate in percentage post challenge test. First 20 family is low resistance group and last 20 families is high resistance group.

**Supplementary Figure S2.**
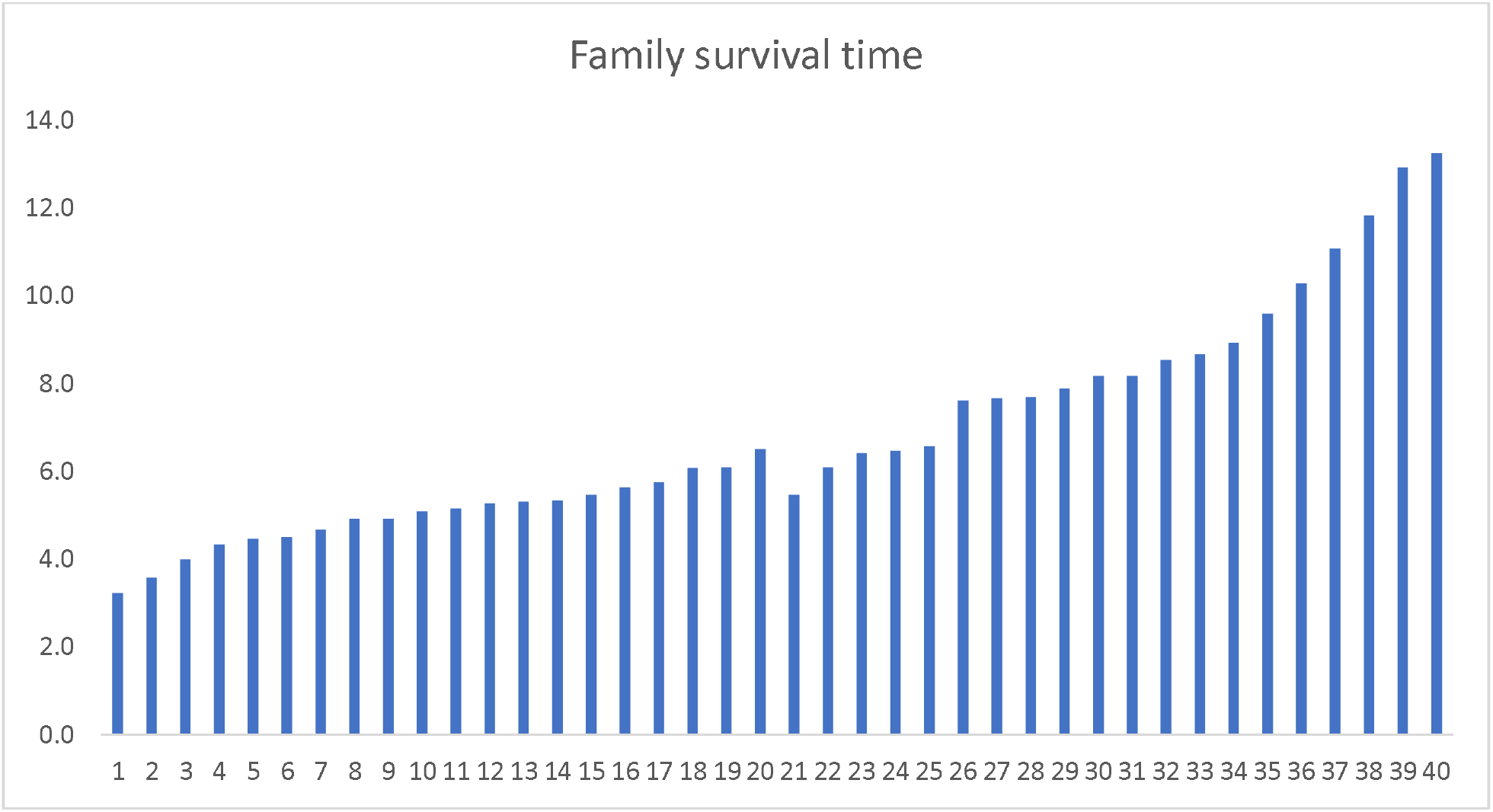
Mean family survival time in day post challenge test. First 20 family is low resistance groups, and last 20 families is high resistance group.

**Supplementary Table S1.** Accuracy between methods using non-imputed or imputed genotypes using 6470 SNPs.

| Trait | Method | Non-imputed |  | Imputed |  | Increase (%) |
| --- | --- | --- | --- | --- | --- | --- |
|  |  | Accuracy±sd<br>(Pearson) | Accuracy±sd<br>(Mathew's) | Accuracy±sd<br>(Pearson) | Accuracy±sd<br>(Mathew's) |  |
| Binary survival | PBLUP | 0.66±0.08 | 0.47±0.11 | - |  | - |
|  | GBLUP | 0.55±0.08 | 0.39±0.12 | 0.65±0.07 | 0.51±0.08 | + 19.0 |
|  | ssGBLUP | 0.55±0.08 | 0.39±0.12 | 0.65±0.07 | 0.51±0.07 | + 19.2 |
|  | BayesR | 0.67±0.08 | 0.55±0.09 | 0.76±0.06 | 0.63±0.09 | + 12.5 |
|  | ML-KAML | 0.67±0.07 | 0.57±0.10 | 0.75±0.06 | 0.63±0.10 | + 12.0 |
|  | DL-MLP* | n.e. | n.e. | 0.71±0.11 | 0.65±0.11 |  |
|  | DL-CNN | n.e. | n.e. | 0.73±0.08 | 0.63±0.12 |  |
| Survival time | PBLUP | 0.65±0.08 | n.e. | - | - | - |
|  | GBLUP | 0.59±0.09 | n.e. | 0.65±0.09 | n.e. | + 10.0 |
|  | ssGBLUP | 0.61±0.07 | n.e. | 0.65±0.09 | n.e. | + 7.1 |
|  | BayesR | 0.70±0.07 | n.e. | 0.75±0.07 | n.e. | + 5.3 |
|  | ML-KAML | 0.69±0.07 | n.e. | 0.75±0.07 | n.e. | + 8.1 |
|  | DL-MLP | n.e. | n.e. | 0.67±0.09 | n.e. |  |
|  | DL-CNN | n.e. | n.e. | 0.70±0.11 | n.e. |  |

n.e. not estimable due to DL-MLP and DL-CNN do not accept un-imputed missing genotype or not estimable for Mathew's coefficient of continuous trait (i.e., survival time). Increase (%) = (accuracy of imputed 6470 SNPs – accuracy of un-imputed 6470 SNPs)/ accuracy of un-imputed 6470 SNPs

**Supplementary Table S2.** Number of SNPs used for each testing set based on its effect with p<0.00001

| Subset | Total SNP used for ssGWAS | ssGWAS SNPs for survival status | ssGWAS SNPs for survival time |
| --- | --- | --- | --- |
| i1t1 | 6470 | 342 | 1448 |
| i1t2 | 6470 | 381 | 1537 |
| i1t3 | 6470 | 356 | 1399 |
| i1t4 | 6470 | 351 | 1561 |
| i1t5 | 6470 | 318 | 1499 |
| i2t1 | 6470 | 335 | 1450 |
| i2t2 | 6470 | 366 | 1598 |
| i2t3 | 6470 | 400 | 1540 |
| i2t4 | 6470 | 318 | 1454 |
| i2t5 | 6470 | 370 | 1401 |
| i3t1 | 6470 | 341 | 1401 |
| i3t2 | 6470 | 365 | 1475 |
| i3t3 | 6470 | 373 | 1514 |
| i3t4 | 6470 | 369 | 1559 |
| i3t5 | 6470 | 361 | 1538 |
| i4t1 | 6470 | 372 | 1576 |
| i4t2 | 6470 | 326 | 1362 |
| i4t3 | 6470 | 357 | 1539 |
| i4t4 | 6470 | 349 | 1383 |
| i4t5 | 6470 | 327 | 1548 |
| i5t1 | 6470 | 329 | 1423 |
| i5t2 | 6470 | 367 | 1480 |
| i5t3 | 6470 | 341 | 1496 |
| i5t4 | 6470 | 362 | 1555 |
| i5t5 | 6470 | 360 | 1502 |
| Mean |  | 353.4 | 1489.5 |
| Min |  | 318 | 1362 |
| Max |  | 400 | 1589 |

## References

Abdollahi-Arpanahi, R., G. Morota, and F. Peñagaricano, 2017 Predicting bull fertility using genomic data and biological information. Journal of Dairy Science 100 (12):9656–9666.

Aguilar, I., I. Misztal, D. Johnson, A. Legarra, S. Tsuruta et al., 2010 Hot topic: A unified approach to utilize phenotypic, full pedigree, and genomic information for genetic evaluation of Holstein final score. Journal of Dairy Science 93 (2):743–752.

Al Kalaldeh, M., J. Gibson, N. Duijvesteijn, H.D. Daetwyler, I. MacLeod et al., 2019 Using imputed whole-genome sequence data to improve the accuracy of genomic prediction for parasite resistance in Australian sheep. Genetics Selection Evolution 51 (1):1–13.

Baker, L.A., M. Momen, K. Chan, N. Bollig, F.B. Lopes et al., 2020 Bayesian and Machine Learning Models for Genomic Prediction of Anterior Cruciate Ligament Rupture in the Canine Model. G3: Genes, genomes, genetics 10 (8):2619–2628.

Bargelloni, L., O. Tassiello, M. Babbucci, S. Ferraresso, R. Franch et al., 2021 Data imputation and machine learning improve association analysis and genomic prediction for resistance to fish photobacteriosis in the gilthead sea bream. Aquaculture Reports 20:100661.

Barría, A., K.A. Christensen, G.M. Yoshida, K. Correa, A. Jedlicki et al., 2018 Genomic predictions and genome-wide association study of resistance against Piscirickettsia salmonis in coho salmon (Oncorhynchus kisutch) using ddRAD sequencing. G3: Genes, genomes, genetics 8 (4):1183–1194.

Benesty, J., J. Chen, Y. Huang, and I. Cohen, 2009 Pearson correlation coefficient, pp. 1–4 in Noise reduction in speech processing. Springer.

Chen, L., C. Li, M. Sargolzaei, and F. Schenkel, 2014 Impact of genotype imputation on the performance of GBLUP and Bayesian methods for genomic prediction. PloS one 9 (7):e101544.

Chicco, D., and G. Jurman, 2020 The advantages of the Matthews correlation coefficient (MCC) over F1 score and accuracy in binary classification evaluation. BMC genomics 21 (1):1–13.

Conesa, A., S. Götz, J.M. García-Gómez, J. Terol, M. Talón et al., 2005 Blast2GO: a universal tool for annotation, visualization and analysis in functional genomics research. Bioinformatics 21 (18):3674–3676.

Crossa, J., Y. Beyene, S. Kassa, P. Pérez, J.M. Hickey et al., 2013 Genomic prediction in maize breeding populations with genotyping-by-sequencing. G3: Genes, genomes, genetics 3 (11):1903–1926.

Daetwyler, H.D., M.P. Calus, R. Pong-Wong, G. de Los Campos, and J.M. Hickey, 2013 Genomic prediction in animals and plants: simulation of data, validation, reporting, and benchmarking. Genetics 193 (2):347–365.

Davey, J.W., and M.L.J.B.i.f.g. Blaxter, 2010 RADSeq: next-generation population genetics. 9 (5-6):416–423.

Dinh Pham, K., J. Ødegård, S. Van Nguyen, H. Magnus Gjøen, and G. Klemetsdal, 2021 Genetic analysis of resistance in Mekong striped catfish (Pangasianodon hypophthalmus) to bacillary necrosis caused by Edwardsiella ictaluri. Journal of fish diseases 44 (2):201–210.

Elshire, R.J., J.C. Glaubitz, Q. Sun, J.A. Poland, K. Kawamoto et al., 2011 A robust, simple genotyping-by-sequencing (GBS) approach for high diversity species. PloS one 6 (5):e19379.

Erbe, M., B. Hayes, L. Matukumalli, S. Goswami, P. Bowman et al., 2012 Improving accuracy of genomic predictions within and between dairy cattle breeds with imputed high-density single nucleotide polymorphism panels. Journal of Dairy Science 95 (7):4114–4129.

Freeman, E.A., and G. Moisen, 2008 PresenceAbsence: An R package for presence absence analysis. Journal of Statistical Software. 23 (11): 31 p.

Gilmour, A., B. Gogel, B. Cullis, S. Welham, R. Thompson et al., 2014 ASReml user guide. Release 4.1 structural specification. VSN International Ltd, Hemel Hempstead, HP1 1ES, UK www.vsni.co.uk.

González-Recio, O., and S. Forni, 2011 Genome-wide prediction of discrete traits using Bayesian regressions and machine learning. Genetics Selection Evolution 43 (1):1–12.

Gowane, G.R., S.H. Lee, S. Clark, N. Moghaddar, H.A. Al-Mamun et al., 2019 Effect of selection and selective genotyping for creation of reference on bias and accuracy of genomic prediction. Journal of Animal Breeding and Genetics 136 (5):390–407.

Gruber, B., P.J. Unmack, O.F. Berry, and A. Georges, 2018 dartr: An r package to facilitate analysis of SNP data generated from reduced representation genome sequencing. Mol Ecol Resour 18 (3):691–699.

Hayes, B.J., P.J. Bowman, A.J. Chamberlain, and M.E. Goddard, 2009 Invited review: Genomic selection in dairy cattle: Progress and challenges. Journal of Dairy Science 92 (2):433–443.

Heidaritabar, M., M.P. Calus, H.J. Megens, A. Vereijken, M.A. Groenen et al., 2016 Accuracy of genomic prediction using imputed whole-genome sequence data in white layers. Journal of Animal Breeding and Genetics 133 (3):167–179.

Henderson, C., 1985 Best linear unbiased prediction of nonadditive genetic merits in noninbred populations. Journal of Animal Science 60 (1):111–117.

Houston, R.D., T.P. Bean, D.J. Macqueen, M.K. Gundappa, Y.H. Jin et al., 2020 Harnessing genomics to fast-track genetic improvement in aquaculture. Nature Reviews Genetics 21 (7):389–409.

Jeong, S., J.-Y. Kim, and N. Kim, 2020 GMStool: GWAS-based marker selection tool for genomic prediction from genomic data. Scientific reports 10 (1):1–12.

Joshi, R., A. Skaarud, M. de Vera, A.T. Alvarez, and J. Ødegård, 2020 Genomic prediction for commercial traits using univariate and multivariate approaches in Nile tilapia (Oreochromis niloticus). Aquaculture 516:734641.

Kilian, A., P. Wenzl, E. Huttner, J. Carling, L. Xia et al., 2012 Diversity arrays technology: a generic genome profiling technology on open platforms, pp. 67–89 in Data production and analysis in population genomics. Springer.

Kim, O.T.P., P.T. Nguyen, E. Shoguchi, K. Hisata, T.T.B. Vo et al., 2018 A draft genome of the striped catfish, Pangasianodon hypophthalmus, for comparative analysis of genes relevant to development and a resource for aquaculture improvement. BMC genomics 19 (1):733.

Li, B., N. Zhang, Y.-G. Wang, A.W. George, A. Reverter et al., 2018 Genomic prediction of breeding values using a subset of SNPs identified by three machine learning methods. Frontiers in genetics 9:237.

Lourenco, D., A. Legarra, S. Tsuruta, Y. Masuda, I. Aguilar et al., 2020 Single-step genomic evaluations from theory to practice: using SNP chips and sequence data in BLUPF90. Genes 11 (7):790.

Luo, Z., Y. Yu, J. Xiang, and F. Li, 2021 Genomic selection using a subset of SNPs identified by genome-wide association analysis for disease resistance traits in aquaculture species. Aquaculture 539:736620.

Masuda, Y., I. Aguilar, S. Tsuruta, and I. Misztal, 2014 Acceleration of Computations in AI REML for Single-step GBLUP Models in Proceedings of the 10th World Congress on Genetics Applied to Livestock Production.

Matthews, B.W., 1975 Comparison of the predicted and observed secondary structure of T4 phage lysozyme. Biochimica et Biophysica Acta (BBA)-Protein Structure 405 (2):442–451.

Meuwissen, T.H., B.J. Hayes, and M.E. Goddard, 2001 Prediction of total genetic value using genome-wide dense marker maps. Genetics 157 (4):1819–1829.

Misztal, I., A. Legarra, and I. Aguilar, 2009 Computing procedures for genetic evaluation including phenotypic, full pedigree, and genomic information. Journal of Dairy Science 92 (9):4648–4655.

Misztal, I., S. Tsuruta, T. Strabel, B. Auvray, T. Druet et al., 2002 BLUPF90 and related programs (BGF90), pp. 743–744 in Proceedings of the 7th world congress on genetics applied to livestock production.

Montesinos-López, O.A., J. Martín-Vallejo, J. Crossa, D. Gianola, C.M. Hernández-Suárez et al., 2019 New deep learning genomic-based prediction model for multiple traits with binary, ordinal, and continuous phenotypes. G3: Genes, genomes, genetics 9 (5):1545–1556.

Montesinos-López, O.A., A. Montesinos-López, J. Crossa, D. Gianola, C.M. Hernández-Suárez et al., 2018 Multi-trait, multi-environment deep learning modeling for genomic-enabled prediction of plant traits. G3: Genes, genomes, genetics 8 (12):3829–3840.

Moser, G., S.H. Lee, B.J. Hayes, M.E. Goddard, N.R. Wray et al., 2015 Simultaneous discovery, estimation and prediction analysis of complex traits using a Bayesian mixture model. PLoS Genet 11 (4):e1004969.

Nani, J.P., F.M. Rezende, and F. Peñagaricano, 2019 Predicting male fertility in dairy cattle using markers with large effect and functional annotation data. BMC genomics 20 (1):258.

Nguyen, N.H., and P.V. Khang, 2021 Genome-Wide Marker Analysis for Traits of Economic Importance in Asian Seabass Lates calcarifer. Journal of Marine Science and Engineering 9 (3):282.

Nguyen, N.H., C. Phuthaworn, and W. Knibb, 2020 Genomic prediction for disease resistance to Hepatopancreatic parvovirus and growth, carcass and quality traits in Banana shrimp Fenneropenaeus merguiensis. Genomics 112 (2):2021–2027.

Nguyen, N.H., P. Rastas, H. Premachandra, and W. Knibb, 2018 First high-density linkage map and single nucleotide polymorphisms significantly associated with traits of economic importance in Yellowtail Kingfish Seriola lalandi. Frontiers in genetics 9:127.

Nguyen, N.P., 2014 Enviromental factors affecting the pathogenesis of Edwardsiella ictaluri in striped catfish Pangasianodon hypophthalmus (Sauvage).

Okeke, U.G., D. Akdemir, I. Rabbi, P. Kulakow, and J.-L. Jannink, 2017 Accuracies of univariate and multivariate genomic prediction models in African cassava. Genetics Selection Evolution 49 (1):1–10.

Palaiokostas, C., 2021 Predicting for disease resistance in aquaculture species using machine learning models. Aquaculture Reports 20:100660.

Pérez-Enciso, M., and L.M. Zingaretti, 2019 A guide on deep learning for complex trait genomic prediction. Genes 10 (7):553.

Pham, K.D., S.V. Nguyen, J. Ødegård, H.M. Gjøen, and G. Klemetsdal, 2020 Case study development of a challenge test against Edwardsiella ictaluri in Mekong striped catfish (Pangasianodon hypophthalmus), for use in breeding: Estimates of the genetic correlation between susceptibilities in replicated tanks. Journal of fish diseases.

Pham, K.D., J. Ødegård, S.V. Nguyen, H.M. Gjøen, and G. Klemetsdal, 2021 Genetic correlations between challenge tested susceptibility to bacillary necrosis, caused by Edwardsiella ictaluri, and growth performance tested survival and harvest body weight in Mekong striped catfish (Pangasianodon hypophthalmus). Journal of fish diseases 44 (2):191–199.

Purcell, S., B. Neale, K. Todd-Brown, L. Thomas, M.A. Ferreira et al., 2007 PLINK: a tool set for whole-genome association and population-based linkage analyses. The American Journal of Human Genetics 81 (3):559–575.

R Core Team, 2015 R: A Language and Environment for Statistical Computing. Vienna, Austria: R Foundation for Statistical Computing; 2014. R Foundation for Statistical Computing. ISBN 3-900051-07-0. http://www.R-project.org.

Robledo, D., C. Palaiokostas, L. Bargelloni, P. Martínez, and R. Houston, 2018 Applications of genotyping by sequencing in aquaculture breeding and genetics. Reviews in aquaculture 10 (3):670–682.

Silva, R.M., J.P. Evenhuis, R.L. Vallejo, G. Gao, K.E. Martin et al., 2019 Whole-genome mapping of quantitative trait loci and accuracy of genomic predictions for resistance to columnaris disease in two rainbow trout breeding populations. Genetics Selection Evolution 51 (1):42.

Sun, J., J.E. Rutkoski, J.A. Poland, J. Crossa, J.L. Jannink et al., 2017 Multitrait, random regression, or simple repeatability model in high-throughput phenotyping data improve genomic prediction for wheat grain yield. The plant genome 10 (2):plantgenome2016.2011.0111.

Tsai, H.-Y., O. Matika, S.M. Edwards, R. Antolín–Sánchez, A. Hamilton et al., 2017 Genotype imputation to improve the cost-efficiency of genomic selection in farmed Atlantic salmon. G3: Genes, genomes, genetics 7 (4):1377–1383.

Tsuruta, S., and I. Misztal, 2006 THRGIBBS1F90 for estimation of variance components with threshold and linear models. Threshold 3 (4).

Van Binsbergen, R., M.P. Calus, M.C. Bink, F.A. van Eeuwijk, C. Schrooten et al., 2015 Genomic prediction using imputed whole-genome sequence data in Holstein Friesian cattle. Genetics Selection Evolution 47 (1):1–13.

Van Sang, N., G. Klemetsdal, J. Ødegård, and H.M. Gjøen, 2012 Genetic parameters of economically important traits recorded at a given age in striped catfish (Pangasianodon hypophthalmus). Aquaculture 344:82–89.

VanRaden, P.M., 2008 Efficient methods to compute genomic predictions. Journal of Dairy Science 91 (11):4414–4423.

VanRaden, P.M., M.E. Tooker, J.R. O’connell, J.B. Cole, and D.M. Bickhart, 2017 Selecting sequence variants to improve genomic predictions for dairy cattle. Genetics Selection Evolution 49 (1):1–12.

Vu, N.T., T.T.T. Ha, V.T.B. Thuy, V.T. Trang, and N.H. Nguyen, 2020 Population Genomic Analyses of Wild and Farmed Striped Catfish Pangasianodon Hypophthalmus in the Lower Mekong River. Journal of Marine Science and Engineering 8 (6):471.

Vu, N.T., N.V. Sang, T.Q. Trong, N.H. Duy, N.T. Dang et al., 2019a Breeding for improved resistance to Edwardsiella ictaluri in striped catfish (Pangasianodon hypophthalmus): Quantitative genetic parameters. Journal of fish diseases 42 (10):1409–1417.

Vũ, N.T., T.Q. TrͰng, L.T. ĐͰnh, H.T.N. Nga, and N.V. HiͰp, 2017 Draft report: Survey results from Project “Investigation for develop national standard regulation: freshwater fish - striped catfish broodtsock and fingerlings - quality requirements”. Draft report to Fishery Department by Research Institute for Aquacultute No 2:25.

Vu, N.T., N. Van Sang, T.H. Phuc, N.T. Vuong, and N.H. Nguyen, 2019b Genetic evaluation of a 15-year selection program for high growth in striped catfish Pangasianodon hypophthalmus. Aquaculture.

Vu, S.V., C. Gondro, N.T. Nguyen, A.R. Gilmour, R. Tearle et al., 2021 Prediction Accuracies of Genomic Selection for Nine Commercially Important Traits in the Portuguese Oyster (Crassostrea angulata) Using DArT-Seq Technology. Genes 12 (2):210.

Whalen, A., G. Gorjanc, and J.M. Hickey, 2020 AlphaFamImpute: high-accuracy imputation in full-sib families from genotype-by-sequencing data. Bioinformatics 36 (15):4369–4371.

Ye, S., J. Li, and Z. Zhang, 2020 Multi-omics-data-assisted genomic feature markers preselection improves the accuracy of genomic prediction. Journal of animal science and biotechnology 11 (1):1–12.

Yin, L., H. Zhang, X. Zhou, X. Yuan, S. Zhao et al., 2020 KAML: improving genomic prediction accuracy of complex traits using machine learning determined parameters. Genome Biology 21 (1):1–22.

Zhu, X., P. Ni, Q. Xing, X. Huang, X. Hu et al., 2021 Genomic prediction of growth traits in scallop using convolutional neural networks. Aquaculture:737171.

Zingaretti, L.M., S.A. Gezan, L.F.V. Ferrão, L.F. Osorio, A. Monfort et al., 2020 Exploring deep learning for complex trait genomic prediction in polyploid outcrossing species. Frontiers in plant science 11:25.

